# Mechanical control of germ cell specification in Arabidopsis anthers

**DOI:** 10.1101/2024.10.17.618875

**Authors:** Chan Liu, Hui Shi, Yuting Han, Pan Wang, Kexin Li, Zhishuai Zhang, Jiazheng Liu, Yafeng Zheng, Linlin Li, Limei Lin, Chen Liang, Binjun Qin, Hua Han, Shunong Bai, Xiao Liu, Wenqian Chen, Feng Zhao

## Abstract

A central question in developmental biology is how the germline is established. We have studied the specification of the male germ cells (GCs) within the anther^1–3^. Hereby, we have focused on the potential role of mechanics, an aspect of anther development which has been very poorly characterized^2,4^. Using a combination of live imaging and mechanical measurements, we provide evidence that GCs originate in a special micro-mechanical niche, where inner tissues exert ‘push’ on outer cell layers, placing them under compression. Mechanical perturbations significantly disrupted the GC specification and patterning. Moreover, we found that the master genetic regulator SPOROCYTELESS/NOZZLE (SPL/NZZ)^5,6^ is central in establishing this micro-mechanical environment by softening the cell wall. The mechanical cues, in turn, stabilize the transcription of *SPL/NZZ*. We propose here an intrinsic growth-derived mechano-chemical feedback loop that drives germ-cell fate acquisition.

## Main

In multicellular organisms, cells interact and coordinate to give special forms with particular functions, accompanied by cell growth and differentiation. Unlike animal cells, plant cells are ‘glued’ together by cell wall and cannot move. As a result, differential growth between neighboring cells or cell clusters leads to mechanical conflicts and stress patterns within tissues during development. These, in turn, can be sensed by cells, guiding their behaviors, such as expansion, division, gene expression and ultimately even fate determination^7^. The link between development and mechanics is often poorly characterized. We addressed this issue in male germ cells (GCs) in higher plants, which are indispensable for maintaining genetic continuity across generations. In most angiosperms, primordial male germ cells, known as archesporial cells (ARs), are specified within the inner cells of anther primordia. As growth and cell division proceed, part of AR descendants is further specified and positioned within the center of anther lobe, surrounded by multiple somatic cell layers. These cells ultimately mature into a population of germ cells, known as pollen mother cells (PMCs), which are destined for meiosis^8,9^ (Fig.1a, Supplementary Fig. 1a). In the past decades, several genetic factors and chemical cues have been identified in guiding the centripetal germ-soma specification^2,4^, however, whether mechanical signals play a role remains unknown.

### Cell wall **(CW)** mechanical heterogeneity parallel with germ-soma segregation

Anther primordia initiate as small radial symmetric domes at the flank of floral meristem. As the four distinct lobes form and elongate around the central filament^10^, the anther shape evolves from a disk to a trapezium, and eventually takes a butterfly-like appearance in cross-sectional view. This transformation is accompanied by cell growth and differentiation (i.e., the specification of germ and somatic cells) ^9,11^ (Supplementary Fig. 1b-c). To evaluate the growth of different cell types, we employed live imaging to track the dynamic development of pre-meiotic anthers, using the cell membrane marker UBQ10:LTI6b-tdTomato to outline the cells (see Materials and Methods). Under our *in vitro* culture condition, we were able to recapitulate five stages of pre-meiotic anther development, matching the well-known morphological and cell differentiation landmarks^9,11^ (Supplementary Fig. 1b-d). Stage 3 marks the morphologically onset of GC lineage specification (Supplementary Fig. 1a). We analyzed the cell area extension in the cross-section plane from stages 3 to 5 and observed a growth gradient from the inner GC lineage to the inner somatic layers and the outermost epidermis (Fig. 1b-c). This growth pattern corresponds to the formation of PMCs, in the inner core of anther lobes, characterized by larger area (in 2D) and volume (in 3D) (Supplementary Fig. 2a-d).

**Figure 1.**
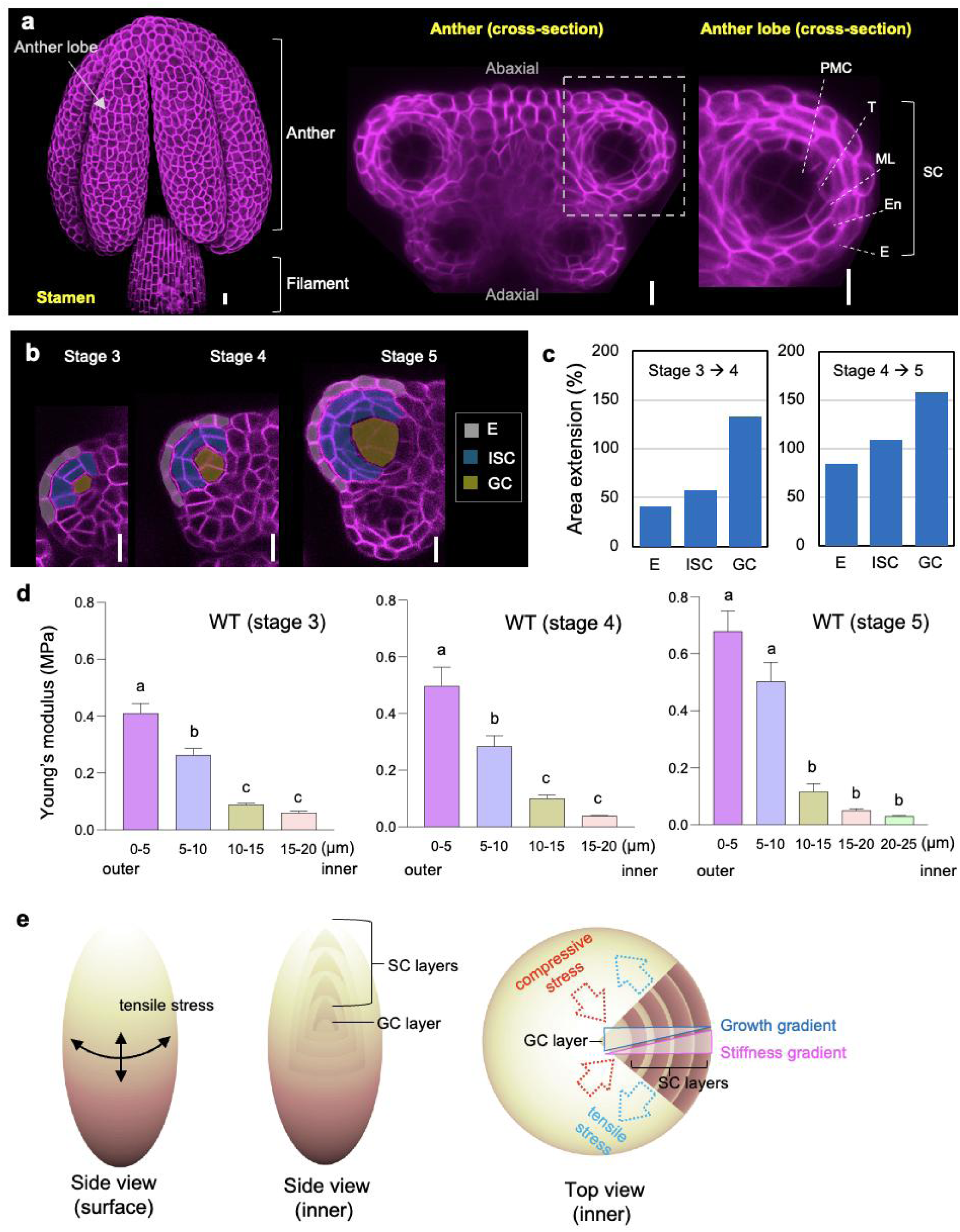
The Arabidopsis male germline initiates in a special micro-mechanical environment. a, Confocal images of Arabidopsis stamen at stage 5 collected through 3D surface reconstructions (left) and transverse optical section (middle) of anther. The right panel is magnified anther lobe from the left white dotted box area showing layered structure in detail. E, epidermis; En, endothecium; ML, middle layer; T, tapetum; PMC, pollen mother cell; SC, somatic cell. b, Time-lapse confocal live imaging records successive anther development from stages 3 to 5. The lineage of different cell clusters is marked in gray (epidermis), blue [inner somatic cell (ISC)], and yellow (germ cell) respectively. c, Relative growth rate of different cell clusters between two successive stages of anthers. Data represents the average value from 5 anther lobes. d, The distribution of cell wall stiffness outside to inside on the transverse plane of anther lobes at stages 3 to 5. Mean values with ±SEM are obtained from three radial directed measurements per lobe with 3 lobes in total for each stage. Different letters on graphs indicate significant difference (*P* < 0.05) calculated by one-way ANOVA followed by Tukey’s test. e, Cartoons showing the tissue stress pattern in the conceptualized anther lobe surface (left panel) and inner cell layers (right panel). Note that there is a growth-derived mechanical conflict between inner germinal and outer somatic cell layers (right panel). Scale bars, 10 μm.

Growth, in essence, is a process of turgor-driven yielding of the CW. According to Lockhart equation, plant cell growth is formulated as: v = φ (P-Y)^12^, where v represents the growth rate, φ denotes CW extensibility, P stands for turgor pressure, and Y is the pressure threshold above which CW yields, allowing growth to occur. The CW resists turgor pressure and controls growth via remodeling its mechanical properties such as CW extensibility and rigidity^13,14^. To understand the mechanical properties of CW in all anther cell layers, we applied atomic force microscopy (AFM) to measure the CW rigidity in cross-sections of anther lobes (see Materials and Methods). We observed that starting from stage 3, a stiffness gradient occurs, with inner tissues being softer than outer cell layers (Fig. 1d stage 3). As the anther development progresses, the outer somatic cells become much stiffer and the stiffness gradient steepens (Fig. 1d stage 4). By stage 5, the outermost two layers exhibit the highest stiffness (Fig. 1d stage 5, Supplementary Fig. 3a). These changes in mechanical heterogeneity correspond with the growth gradient across different cell layers (Fig. 1c), and coincide with germ-soma differentiation during early anther morphogenesis.

### Mechanical conflicts in pre-meiotic anther lobes

The pattern of stiffness intensity echoes the epidermal theory of growth which has been validated for many aerial plant organs (e.g., leaves, sepals, stems, immature seeds etc.)^15–19^, showing the epidermis is under tension and inner tissues are under compression (Fig. 1e). We next examined the directions of the stresses at play. There is compelling evidence, that cortical microtubules (CMTs) align along the main direction of stress within cells, and multicellular tissues^15,16^. Their distribution can, therefore, be used to explore stress patterns. We examined CMT distribution dynamics across different layers during anther development from stages 3 to 5. At stage 3, the CMTs are more anisotropically aligned on the surface of the primordia, predominantly in the circumferential direction (Fig. 2a, e). This CMT distribution pattern is very similar to what has been observed in young leaf primordia, early hypocotyl, stem, and young seeds, which could be explained as CMTs aligning along the maximal tensile stress on a pressurized ellipsoid or cylindrical surface^15–19^ (Fig. 1e). However, as anther primordia develop, the CMTs at the surface become less organized, while their alignment, (expressed as microtubule anisotropy) gradually increases in the inner layers (Fig. 2b-g). The CMT signal intensity in GCs is very weak, with a more isotropic distribution (Fig. 2a-g). These observations suggest a different patterns of stress across layers. As GCs enlarge (Fig. 1b-c; Supplementary Fig. 1c) and also the outer somatic cell layers stiffen (Fig. 1d), a strong mechanical heterogeneity at the interface between GCs and adjacent cells seems to appear.

**Figure 2.**
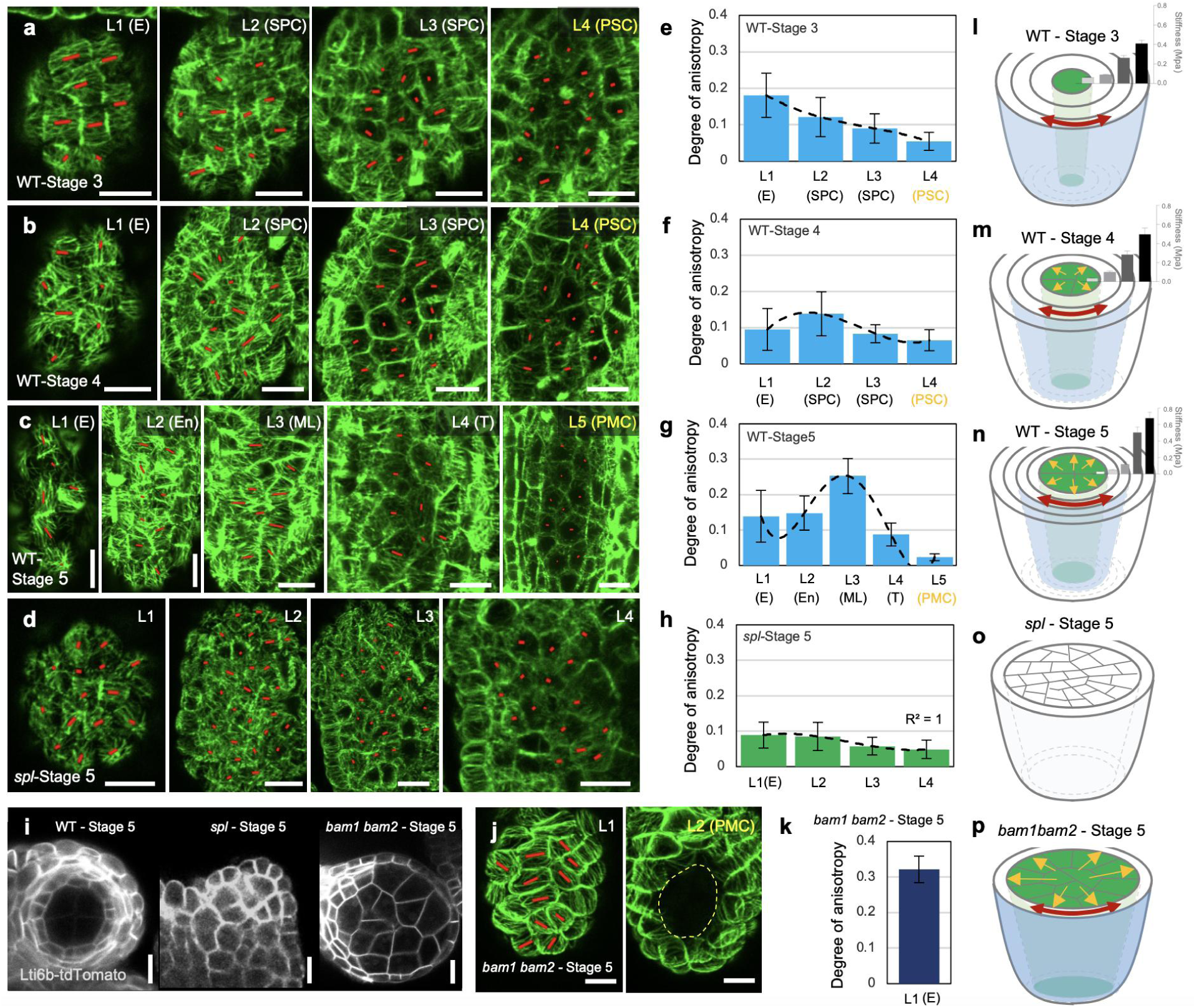
Cortical microtubules (CMTs) organization in WT, *spl* and *bam1 bam2* mutants. a-d, CMTs distribution of 35S:GFP-MBD in different anther cell layer of WT at stage 3 (a), stage 4 (b) and stage 5 (c), and *spl* at stage 5 (d). The orientation and length of red bars represent average microtubule orientation and degree of anisotropy in a single cell computed with FibrilTool. L, layer; E, epidermis; SPC, secondary parietal cell; PSC, primary sporogenous cell. e-h, Quantification of CMTs anisotropy in WT (e-g, n = 46 cells for stage 3, n = 48 cells for stage 4, and n = 42 cells for stage 5) and *spl* (h, n = 83 cells) anthers. i, Cross optical section of UBQ10:LTI6b-tdTomato (LTI6b-tdTomato) labeled WT, *spl*, and *bam1 bam2* anther lobes. j-k, CMTs distribution of 35S:GFP-MBD in *bam1 bam2* (j) and its quantification of anisotropy degree (k, n = 11 cells). Similar CMTs distributions in WT, *spl*, and *bam1 bam2* were observed from at least 3 different anthers. l-p, Schematic representation of CMTs organization, cell wall stiffness, and predicted tension distribution in variant anther lobes. Green cells represent GCs; black-gray-white bars in (l-n) represent the stiffness gradients in stages 3-5 WT anther lobes; yellow arrows indicate the pushing from the GCs; light blue surface indicate the layer in which the degree of CMTs anisotropy is highest; Accordingly, the red double-headed arrows indicate the cell layers which are supposed to be apparently stretched. scale bars, 10 μm.

To further test whether this specific CMT alignment pattern is indeed relevant to the growth-derived pushing force from GCs, we firstly examined *spl*, in which there is no GCs in the lobe^5,6^ (Supplementary Fig. 2e). We assumed that if the better alignment of CMTs is indeed correlated with GC enlargement, such a pattern would be absent in *spl*. Coincide with the absence of enlarged GC in the inner anther lobes, (Supplementary Fig. 2e-f), we did not observe very anisotropic CMTs alignment in either epidermis or inner cell layers of *spl* anthers at stage 5 (Fig. 2d, h). This uniform and isotropic CMTs distribution is consistent with the more rounded and smaller anther shape seen in *spl* (Supplementary Fig. 4), suggesting that the GC enlargement might also contribute to overall anther growth and shape.

Secondly, we investigated CMT alignment in *bam1bam2* anthers. In contrast with *spl*, where there is no GCs, the inner somatic cells of *bam1bam2* anthers are all replaced by GCs, resulting in direct interaction between the epidermis and GCs^20^ (Fig. 2i). We hypothesized that if the shift in anisotropic CMT alignment toward the inner cell layers (Fig. 2a-g) is relevant to the increase of somatic cell layers, then in *bam1bam2*, where the epidermis is the only layer subjected to the tension from GCs, we should observe strong CMTs alignment on the *bam1bam2* anther surface. As expected, while the CMT signals in the GCs of *bam1bam2* anthers were too weak to quantify, we observed highly anisotropic CMT alignment along the circumferential direction on the surface of stage 5 *bam1bam2* anthers (Fig. 2j-k and Supplementary Fig. 5). Altogether, the results above indicate that during the early anther development, there exists a mechanical conflict between germ and soma. This conflict may originate from the stiffness gradient between somatic and germinal regions (Fig.1d-e; Fig. 2l-p).

### Mechanical perturbations influence the germ cell specification

Given the GCs reside in such a very special mechanical environment, we hypothesized that mechanical signals might play a role in GC specification. As previously noted, the mechanical conflicts originate from the differential growth between germinal and somatic regions. To test this hypothesis, we performed two types of experiments to tune the growth parameters and observed their effects on GC specification. We first used the GFP-AGO9 as a germinal lineage marker^21^ to quantify the dynamics of GC specification. Taking advantage of live imaging, we found AGO9 signals in anther primordia were activated starting from stage 3 in two files of subepidermal layers, which corresponds to SPCs (i.e., two descendants given by AR division). Over about 4 days’ culture *in vitro*, the AGO9 proteins gradually localized to the GC lineage, eventually reaching a relatively stable GC (i.e., PMC) number before meiosis (at anther stage around 5 and/or 6) (Supplementary Fig. 6). This AGO9 expression pattern aligns with the proposed lineage model^8^ (Supplementary Fig. 1a) and reinforces the notion that anther stage 3 is an appropriate starting point for studying GC specification.

Using this tracking system, we first applied osmotic treatment by adding mannitol and sorbitol (to exclude the protentional artifacts caused by mannitol signaling transduction^22^) in the culture medium to reduce the tissue turgor pressure. We assumed this treatment would lower osmolarity, primarily affecting the faster-growing cells, and thereby minimize mechanical conflicts between the anther cell layers. Specifically, we monitored the anther growth and GC specification starting from stage 3 under a mild osmotic treatment condition (see Materials and Methods, Supplementary Fig. 7a-f). Since the anther growth is reduced on mannitol and sorbitol medium, we prolonged the culture time to allow sufficient growth for the anthers to reach a typical butterfly-like shape (around developmental stage 5/6), facilitating an equivalent comparison with the control group (Supplementary Fig. 7a-f). Under these conditions, stamens could differentiate normally into anther and filament structures, while the anthers appeared narrower and smaller (Fig. 3a-k; Supplementary Fig. 7a-f). Importantly, the number of AGO9-labeled GCs significantly decreased, with some extreme cases (13 out of 26, 50% locules for mannitol treatment; and 5 out of 21, 24% locules for sorbitol treatment) showing no GCs in the anther lobes (Fig. 3d-k, q-r). These results indicate that turgor-driven growth is essential for GC specification.

**Fig. 3.**
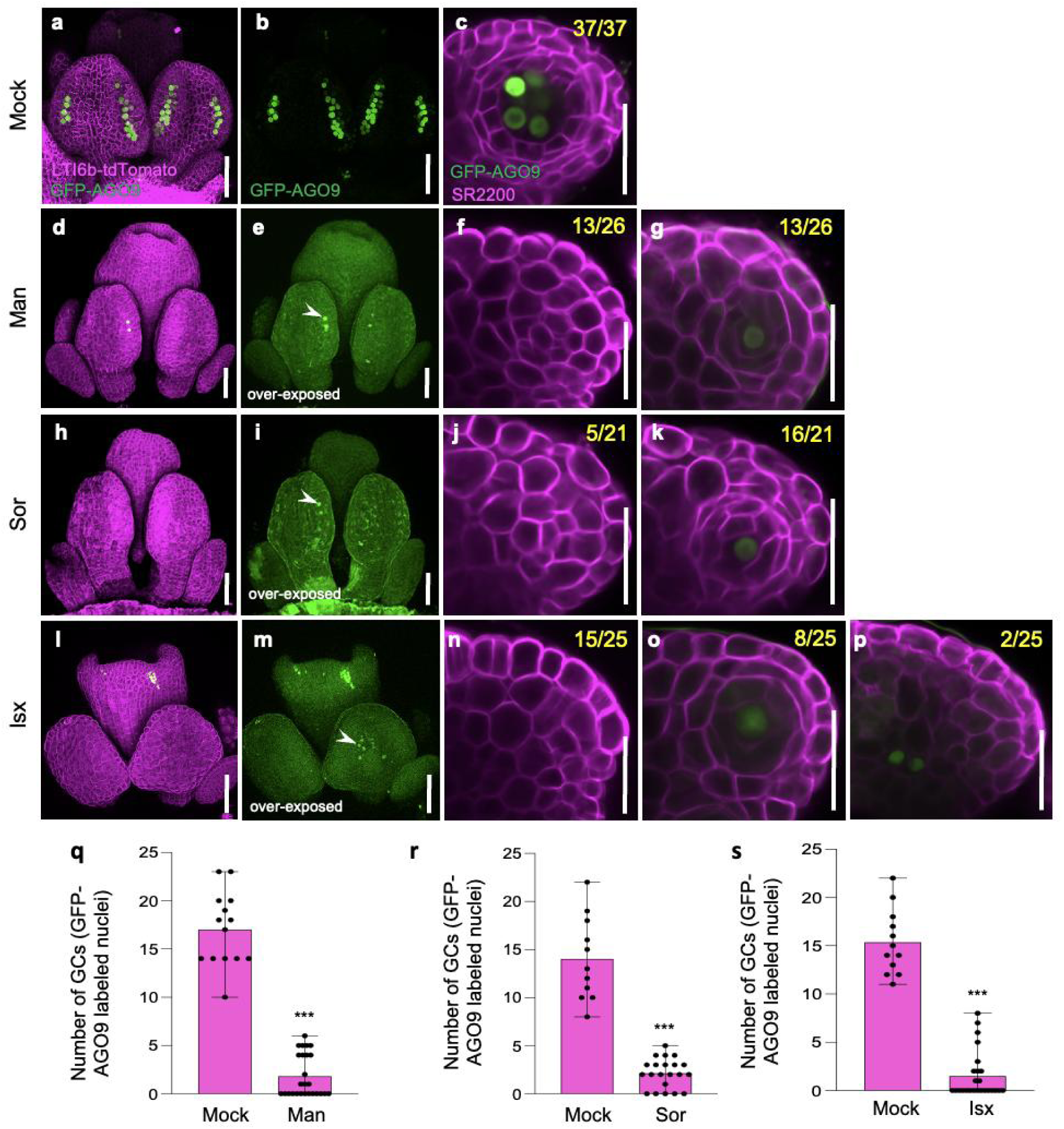
Mechanical perturbations significantly affect germ cell specification. a-p, Confocal images of anthers [from LTI6b-tdTomato proAGO9:EGFP-AGO9 (GFP-AGO9) plants] collected from the 3D reconstruction (the left two columns) and cross-optical sections (the other pictures) that untreated (a-c) or treated with mannitol (Man) (d-g), sorbitol (Sor) (h-k), and isoxaben (Isx) (l-p). Arrowheads indicate the AGO9 signals. Yellow numbers present the statistics of individual situation. q-s, Quantification of GC number formation under different treatment (37 locules were analyzed for mock; 26, 21, 25 locules were analyzed for mannitol, sorbitol, and isoxaben treatments, respectively). Values are mean ±SD. *** Student’s *t*-test *P* < 0.001. Scale bars, 50 µm for 3D reconstruction images, 25 µm for cross-optical section images.

We previously indicated that the mechanical conflicts between the germ and soma layers likely originate from the stiffness gradient between outer and inner tissues (Fig. 1d). Among all wall components, cellulose is the most rigid and serves as the primary load-bearing element in the primary CW in most organs. Blocking the cellulose biosynthesis, in principle could soften the CW. Given that the CWs of outer cell layers are stiffer than those of the inner layers (Fig. 1d), we assumed that reducing wall stiffness would have a greater impact on the outer hard walls, thereby minimizing spatial differences in wall stiffness. Following this idea, we performed a second experiment–treating the anther primordia with isoxaben (Isx), a cellulose biosynthesis inhibitor (see Materials and Methods, Supplementary Fig. 7g-h). Under these conditions, 60% (15 out of 25) of the locules produced no GCs at all (Fig. 3l-p, s). In the remaining cases, GC specification was significantly perturbed (Fig. 3l-p, s). Surprisingly, GCs can even occasionally be formed close to the vascular tissues (2 out of 25, 8%) (Fig. 3p). These findings suggest that CW-mediated mechanical cues participate in the guidance of GC specification and patterning.

### SPOROCYTELESS mediates mechano-chemical feedback in GC specification

The results above indicate that mechanical cues are involved in guiding GC specification. We then ask how is the micro-mechanical environment established? We have proposed that the rapid growth of the inner core of anther lobes is the driving force behind mechanical conflicts. When *SPL/NZZ*, a master gene gating the GC specification^5,6^, is mutated, the GC formation is blocked and mechanical conflicts appear to vanish (Fig. 2d, h and o; Supplementary Fig. 2e-f). To further explore this, we quantified wall stiffness distributions in the cross-sections of *spl* anthers from stages 3 to 5. As expected, we found that the wall mechanical heterogeneity in the anther lobes was eliminated (Fig. 4a; Supplementary Fig. 3b), suggesting that SPL is involved in mechanical regulation. To test whether SPL is indeed required to establish the micro-mechanical environment (i.e., a mechanical gradient) in the anther lobe, we applied *ag-1* AGGR inducible system to spatial-temporally activate *SPL* expression and observed the resulting changes in the CW stiffness. In *ag-1* AGGR, the stamens are replaced by petals. After DEX induction, *AG* activates *SPL* expression at the lateral margin of petals, thereby locally triggering microsporocyte formation^23^ (Fig. 4b; Supplementary Fig. 8a-c). We successively tracked this process by confocal imaging and found 2 days after induction, GCs are recognizable by their large size and AGO9 expression (Fig. 4c, Supplementary Fig. 8d-e). We then applied AFM to measure the wall stiffness following *SPL* activation. We found, that starting from 2 days after induction, the cell walls from the inner side of petal margins appeared softer and established a stiffness gradient. This gradient sharpened as the petals develop (Fig. 4d; Supplementary Fig. 9a). For further confirmation, we took advantage of the Brillion microscope, by which we could probe the viscoelastic property of live plant tissues in a non-destructive and contact-free manner^24–26^. We also found a softening at the margin of petals after the activation of *SPL/NZZ* (Supplementary Fig. 9b-g). These results indicate that SPL indeed could promote the formation of a micro-mechanical environment by softening the local cell wall stiffness.

**Fig. 4.**
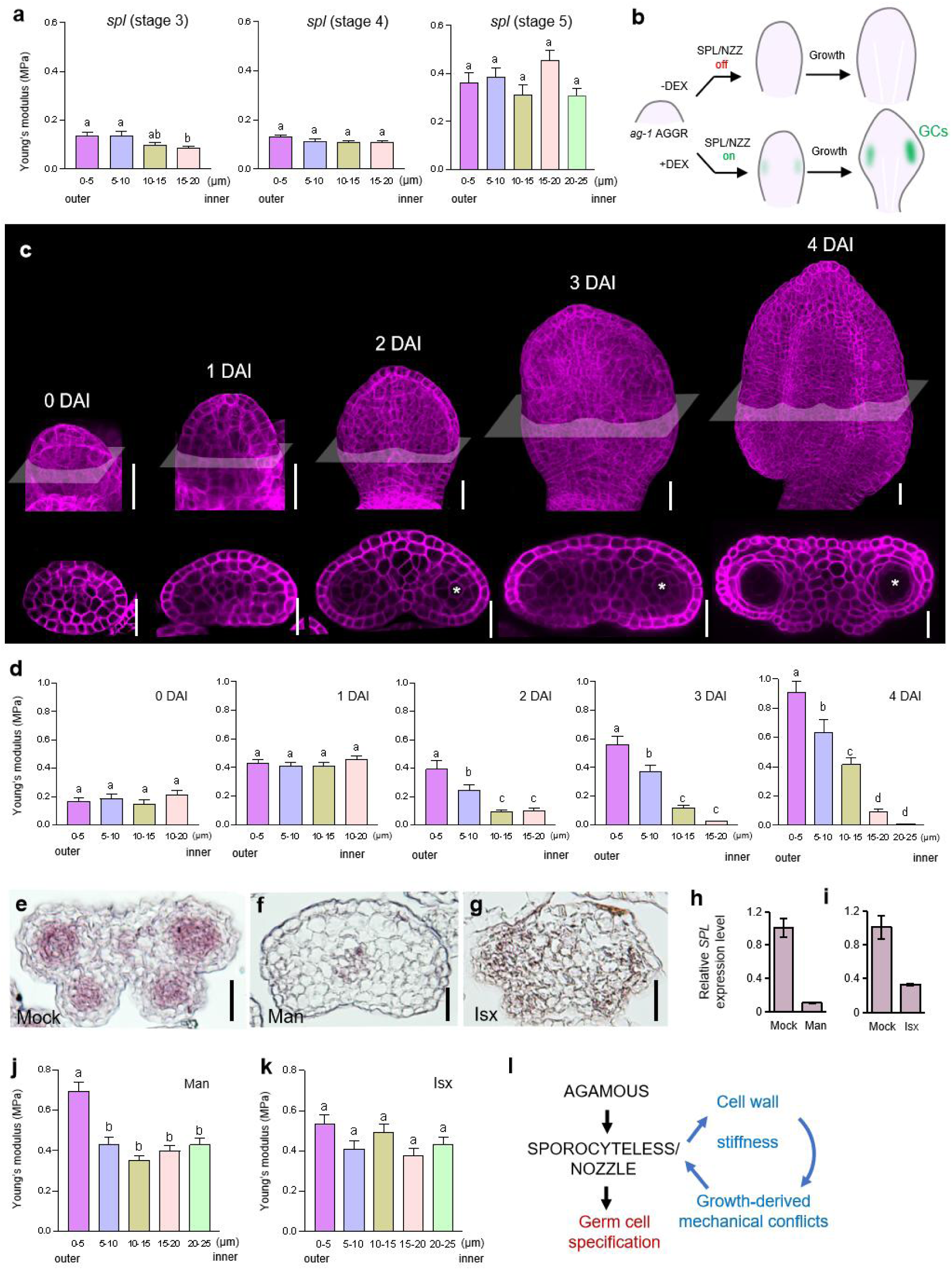
A SPL/NZZ-mediated mechano-chemical feedback drive the male germline specification. a, Cell wall stiffness patterning, represented by Young’s modulus in the cross-section of *spl* anther at stage 3-5. Values are mean ±SEM. b, Schematic representation of the *ag-1* 35S:AG-GR (*ag-1* AGGR) inducible system. C, Detailed images showing GC inductions in *ag-1* AGGR petals. Upper panel, maximum projection of confocal images on petals at different induction time; lower panel, optical cross-section of petals after single DEX induction. DAI, day after induction. White asterisks indicate GC. d, Evolution of cell wall stiffness patterning in the petal of *ag-1* AGGR after single DEX induction. Values are mean ±SEM. Different letters on graphs indicate significant difference (*P* < 0.05) calculated by one-way ANOVA followed by Tukey’s test. e-g, Patterns of *SPL* transcript accumulation in transverse sections through untreated (e), mannitol-treatment (f), and isoxaben-treatment (g). h-i, Quantification of *SPL* transcripts after mechanical perturbations. The graphs show one representative assay of replicates. Bars represent means and error bars indicate ±SD. A *t-*test was used to compare *SPL* expression level in mock and treatments; *** indicate *P* < 0.001. j-k, Cell wall stiffness patterning in anther lobes after mannitol (j) and isoxaben (k) treatment. Values are mean ±SEM. For AFM measurements in (a, d, j and k), three radial-directed linear regions from outside to inside (0-20/25 μm) of three anther lobes were recorded. A SPL-mediated mechano-chemical feedback loop in regulating GC fate determination. Scale bars, 20 μm (c), 25 μm (e-g).

We previously demonstrated that mechanical perturbations interfere with GC specification (Fig. 3). Building on this, we applied osmotic, and Isx treatment in the *ag-1* AGGR system, and found these mechanical interventions, which hinder growth, also significantly impeded GC specification in homeotic petals (Supplementary Fig. 10). These results suggest that while activated by AG, SPL on its own is not enough to induce GC fate without an appropriate mechanical niche. To further investigate whether mechanical regulation of GC specification depends on SPL, we performed in situ hybridization assay and quantitative real-time polymerase chain reaction (qPCR) to detect the *SPL* transcripts following the mechano-perturbations. We found that after mechanical perturbations, accompanied by the abolishment of mechanical gradient, the transcription of *SPL* was significantly repressed in the anther locules (Fig. 4e-k). Altogether, these results reveal that the micro-mechanical environment is crucial for maintaining *SPL* transcription, which in turn safeguards the GC specification.

In conclusion, our results indicate that, although the GC specification is regulated by genetic cues, mechanical signals are also essential. Supporting this, a recent study demonstrated that the mechanical interactions between tissue layers underlie anther morphogenesis^27^. Our findings further indicate that the growth-derived mechanical conflict is at least in part driven by SPL/NZZ-mediated cell wall softening. Moreover, the well-established micro-mechanical niche helps preserve *SPL/NZZ* transcription, therefore safeguarding the GC specification (Fig. 4l). In animals, mechanical cues have been known to play a significant role in guiding stem cell status, cell differentiation, and/or germ-cell fate determination^28–30^. Our results suggest that this might also be the case in plants.

## Materials and Methods

### Plant materials and growth conditions

Arabidopsis (*Arabidopsis thaliana*) ecotype Columbia-0 (Col), mutants and transgenic plants in the Col or Landsberg erecta (Ler) background were used in this study. The mutants, including *spl-2*^31^, *bam1 bam2*^32^, and the transgenic lines containing pro35S:GFP-MBD^33^, proUBQ10:LTI6b-tdTomato^34^, proAGO9:EGFP-AGO9^35^ and *ag-1* pro35S:AG-GR (*ag-1* AGGR)^36^, all have been described previously. The seeds were sterilized, sown onto Murashige and Skoog (MS) medium for germination, and cultured in vitro. After one week, the seedlings were transplanted to soil and grown under long-day conditions (16-h light/8-h dark; 170 μEs^-1^m^-2^ light intensity provided by a mixture of cool [3,000 K] and warm [6,500 K] white LED lamps) at 22°C with 60% humidity. For the living imaging of anthers, the needless flower buds were dissected away carefully, and the left whole inflorescences were cultured *in vitro* on apex culture medium (ACM) until each acquisition.

### Atomic Force Microscopy (AFM) imaging and data processing

To investigate the mechanical properties of internal anther tissues, we employed cryo-sectioning for sample preparation. In brief, inflorescences were rinsed three times with pre-cooled distilled water, followed by gently blotting the surface with absorbent paper to remove excess moisture. To avoid the potential influence of fixatives on changing the cell wall structure and properties, we soaked the tissues directly in the ice-cold 75% OCT embedding solution for 10 min under vacuum to ensure full infiltration of the OCT into the tissue spaces. Afterward, the samples were embedded in pure OCT and stored at -80℃. For sectioning, the embedding blocks were retrieved in a Leica frozen microtome (CM1950) at -20℃ for 30 min. Sections of 20 µm thickness were then prepared.

AFM experiments were performed using Bruker Nanowizard V coupled to an inverted Zeiss observer 3 microscope, driven by NanoWizard SPM Release software version 8.0. The acquisitions were carried out using the QI (Quantitative Imaging) Mode. All measurements were carried out under water at room temperature to prohibit cell plasmolysis. We used a tip with a nominal radius of 8 nm (RFESPG-75 model; Bruker) mounted on a silicon cantilever with a nominal force constant of 3 N/m. The probe’s spring constant was calibrated before each measurement. Scan size was generally from 100×100 μm^2^ to 250×250 μm^2^ with a resolution of 128×128 pixels depending on the sample size under hybrid stage mode. The applied force trigger was 60 nN, a force corresponding to an indentation of 250-1500 nm.

Data analysis was done using JPK Data Processing Nanoscope Analysis Version 8.1, Sneddon model was used for parameter fitting, cone was selected for Tip shape and 19 deg was selected for Half-Cone Angle. At the same time, frame selection method was also used for data analysis, and frame selection was conducted successively from the anther epidermis to the inner tissues. The size of each frame (square) was about 5µm and contained 16-36 force curves, corresponding to a single cell area. Data getting from three lines of squares along a radial direction in a single anther lobe were collected for statistic analyzation. At least 12 anthers from 3 inflorescences we examined for each period of each sample type.

### Fixing and clearing of plant tissue

Inflorescences was fixed and cleared prior to microscopy imaging according to the previous procedure^37^. Briefly, Samples were first fixed in a 4% (w/v) paraformaldehyde (Sangon Biotech) under vacuum for 10 min, and then incubated at 4°C overnight. Afterward, the fixative was removed and samples were washed three times with 1× PBS. The samples were then incubated in ClearSee solution (10% xylitol (w/v), 15% sodium deoxycholate (w/v), and 25% urea (w/v) in water) in the dark for one week. One day prior to microscopy, samples were stained with SCRI Renaissance 2200 (SR2200) for 1 h in the dark, washed, and mounted on the ACM for imaging.

### Confocal laser scanning microscopy

All confocal images were captured using Nikon AXR laser scanning microscope with a 40× water-immersion objective. GFP was excited with a 488 nm argon laser and emission collected from 500 to 550 nm. tdTomato were excited with a 561 nm argon laser and emission collected from 570 to 620 nm. Cell wall stained with SR2200 was excited with a 405 nm argon laser and emission collected from 425 to 475 nm. All quantitative analyses were done with Fiji software.

### In situ hybridization and RT-qPCR analysis

RNA in situ hybridization experiments were performed as previously described^38^. Briefly, inflorescences in good condition were fixed in 4% (w/v) paraformaldehyde and kept on ice for 6-12 h. Following the dehydration (on ice) and clearing in Histo-Clear (Sangon Biotech), the samples were embedded into paraplast (Sigma-Aidrich) for sectioning. sections of 8-µm were cut and dewaxed before being treated with proteinase K (1µg/ml) for 30 min. Digoxigenin-labeled RNA probes were subsequently hybridized to the slides. To synthesize RNA probes, specific cDNA fragments of target genes were amplified using PCR with primers containing T7 promoter sequences (Supplementary table 1). Details of *in vitro* RNA probe synthesis, hybridization and signal detection can be found at http://www.its.caltech. edu/∼plantlab/protocols/insitu.html.

For RT-qPCR analysis, total RNAs were extracted from Col-0 inflorescences under mannitol, isoxaben treatments or without any treatment, which were cultured on ACM for 4 days, using a RNeasy Plant Mini Kit (Omego). cDNAs were synthesized using reverse transcriptase according to the manufacturer’s instructions (Seven) using oligo(dT) primers (TSINGKE). Primer sequences are shown in Supplementary table 1.

### Cell volume calculation

The 3D cell alignment, denoise, and instance segmentation pipeline (3DCADS) is a deep learning-based framework designed for cell instance segmentation of electron microscopy (EM) sequence images^39^. It provides accurate and robust 3D cell reconstruction with high computational efficiency. The first stage involves the registration of EM sequence images. The second stage focuses on image enhancement and denoising. In the third stage, a Transformer-based neural network is employed to perform semantic segmentation. The fourth stage utilizes a super voxel-based clustering algorithm for cell instantiation. Finally, the morphological parameters of the cell instances were analyzed and statistically evaluated, as well as the topological connectivity was visualized at the fifth stage. Using this framework, we successfully reconstructed the anther of Arabidopsis, obtaining its topological connectivity network, and analyzed the morphological parameters and volume of the distribution of different cell types.

### Brillouin microscopy assay

The Brillouin microscopy setup was described previously^40^. Briefly, to acquire Brillouin spectra, the laser beam was focused into the center of sample that channel by an objective lens (OBJ, NA = 1.2, magnification: 60×). The backward scattered light was collected by the same OBJ, reflected at the polarized beam splitter (PBS), and coupled to the single-mode fiber (SMF). The spectral pattern is imaged by the CMOS camera (iXon, Andor) after relating through a 4f system consisting of spherical lenses and spatial filter, the spectral pattern is imaged by the CMOS camera (ORCA-Fusion, Hamamatsu). A Matlab software was developed to drive both the stage and the camera simultaneously and output the Brillouin frequency shift data by automatically least square Lorentzian fitting the Brillouin spectra. The Brillouin measurement path, indicated by the dashed orange arrow in the figures, follows a medial-to-margin direction in petals, with 100 measurement points collected at 1 micron interval. The Brillouin frequency shifts are calculated as follows:

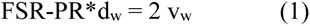

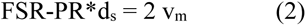

where d_w_ are d_s_ are the peak distance (d) of water (w) and sample (s), FSR (Free Spectral Range) is 30 GHz, v_w_ (Brillouin frequency shift of water) is 7.5 GHz. Firstly, the dw was measured and then was pluged into equation (1) to get the PR (pixel range), which is substituted into equation (2) and together with the measured d_s_ to calculate the Brillouin frequency shift of measured sample (v_m_).

### Alexander Staining Assay

Inflorescences from *ag-1* AGGR plants, either treated or untreated with DEX, were soaked in FAA solution (ethanol: acetic acid = 3:1) for fixation overnight at 4℃. The Petals were then dissected from the inflorescences and mounted on microscope slides in a drop of Alexander’s staining solution. After incubation for 1 h in a wet box at 56℃, the staining solution was rinsed off with distilled water several times. Images were taken using a Zeiss stereomicroscope (Imager. M2).

### Chemical solutions and treatments

Dimethyl sulfoxide (DMSO) was used as a solvent for dissolving isoxaben. For treatment with isoxaben, wild-type (WT) inflorescences inserted onto the ACM were incubated with isoxaben (400 nM, Sigma-Aidrich) for 1 h. For mannitol (0.05 M, Sangon Biotech) and sorbitol (0.1 M, Sangon Biotech) treatments, the powder was directly added to the ACM when autoclaving. Ethanol was used to dissolve dexamethasone (DEX, 10 mM, Macklin). For a single DEX treatment, inflorescences were dipped in water containing 10 µM DEX and 0.05% Silwet L-77 for 2 min. Anthers cultured on the ACM for 8 days after the treatment were collected for phenotype characterization and quantification. Anthers cultured for 4 days after the treatment were collected for qPCR and AFM test.

For the *ag-1* AGGR system, the inflorescences were inserted on the ACM 30 min after DEX treatment, and then these inflorescences underwent isoxaben (600 nM) treatment as mentioned above. For mannitol and sorbitol treatments, inflorescences growth on the ACM 1 d after DEX treatment, and then these inflorescences were moved to ACM plates containing mannitol (0.05 M) or sorbitol (0.1 M).

### Statistical analysis

Statistical analyses were performed in Microsoft Excel or GraphPad Prism. One-way analysis of variance (ANOVA) used for multiple comparisons with a post-hoc Tukey test.

## Acknowledgements

We would like to thank Olivier Hamant and Jan Traas for critical reading of the manuscript and providing valuable suggestions. We would also like to thank Yuchen Long for discussion and technical supports on AFM measurement and data analysis. We thank Ruben Gutzat and Ortrun Mittelsten Scheid for sharing us with the GC marker line proAGO9: EGFP-AGO9; Toshiro Ito, Genji Qin and Xiaoping Gou for their generous gift for the seeds of *ag-1* AGGR, *spl-2* and *bam1 bam2* respectively. This work was funded by the National Natural Science Foundation of China (32370369), Basic Research Field Project “Science and Technology Innovation Action Plan” of the Shanghai Science and Technology Commission (22JC1402800), the Fundamental Research Founds for the Central University, Shaanxi Basic Research Project for Chemistry (22JHQ057), the Youth Program of National Natural Science Foundation of China (32400570), and the Postdoctoral Science Foundation of China (2024M754214) and Shaanxi Province (2023BSHYDZZ58).

## Author contributions

F.Z. and W.C. conceived the project and designed the research. C.L. carried out most of the experiments.

H.S. carried out AFM and in situ hybridization experiments. Y.H. and P.W. generated genetic materials; carried out chemical treatments, participated in the phenotypic analysis and RT-qPCR analysis. C.L. and B.Q. contributed to the data analysis. K.L. and Z.Z. performed Brillouin microscopy assay under the supervision of X.L.. J.L., Y.Z., L.L., and L.L. produced the EM data and conducted cell volume calculation under the supervision of H.H. and S.B.. F.Z., W.C., and C.L. analyzed the data and wrote the manuscript with input from all authors.

## Additional inforamtion

Supplementary figures and table 1

## Supplementary figures and table 1

**Supplementary Fig. 1.**
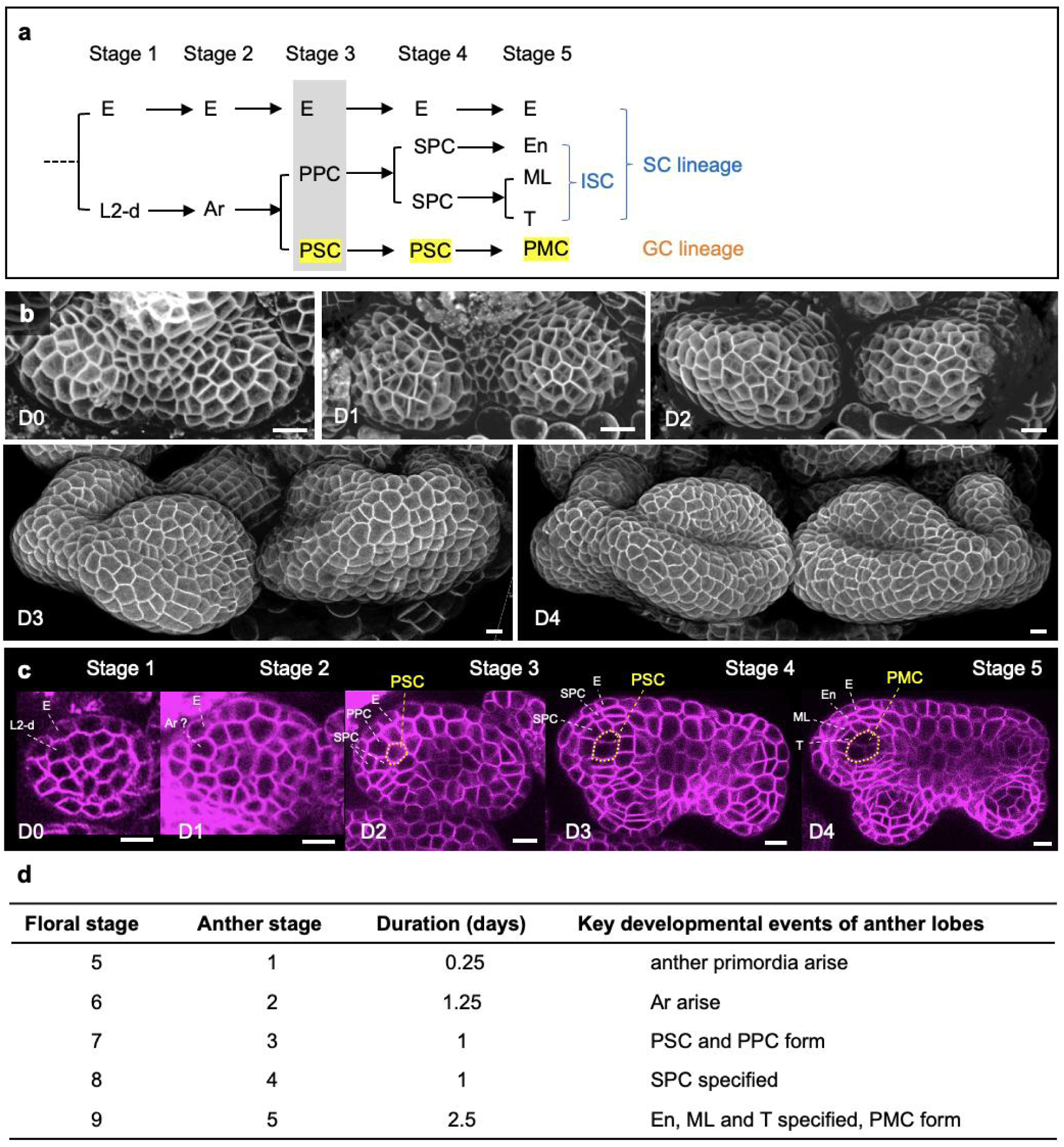
The developmental process of Arabidopsis anther. (a) A schematic model of cell lineage in pre-meiotic anther development. Specifically, the SC lineage and GC lineage are mainly L2 layer-derived (L2-d) at stage 1 except for epidermis. At stage 2, some cells in the L2 layer become archesporial cells (Ar), which divide to produce the PPC (primary parietal cell) and the PSC (primary sporogenous cell) at stage 3. Then the PPC divide to form two layers of SPC (secondary parietal cell) at stage 4. Subsequently, the outer SPC form the endothecium (En), and inner SPC form the tapetum (T) and the middle layer (ML). At the same time, the inner most PSC give rise to the PMC (pollen mother cell) at stage 5. (b) 3D reconstruction of confocal live imaging showing the succussive development of pre-meiotic anthers, which is cultured in vitro for 4 days. D, day. (c) Representative optical cross-sections of anthers at different developmental stages that correspond to different culture days. Cell lineage for different cell types could be tracked in plane. Dashed yellow lines show the GC lineage. (d) Summary of the corresponding of floral stages (5-9) with anther stages (1-5), key developmental events, and duration time for each stage.

**Supplementary Fig. 2.**
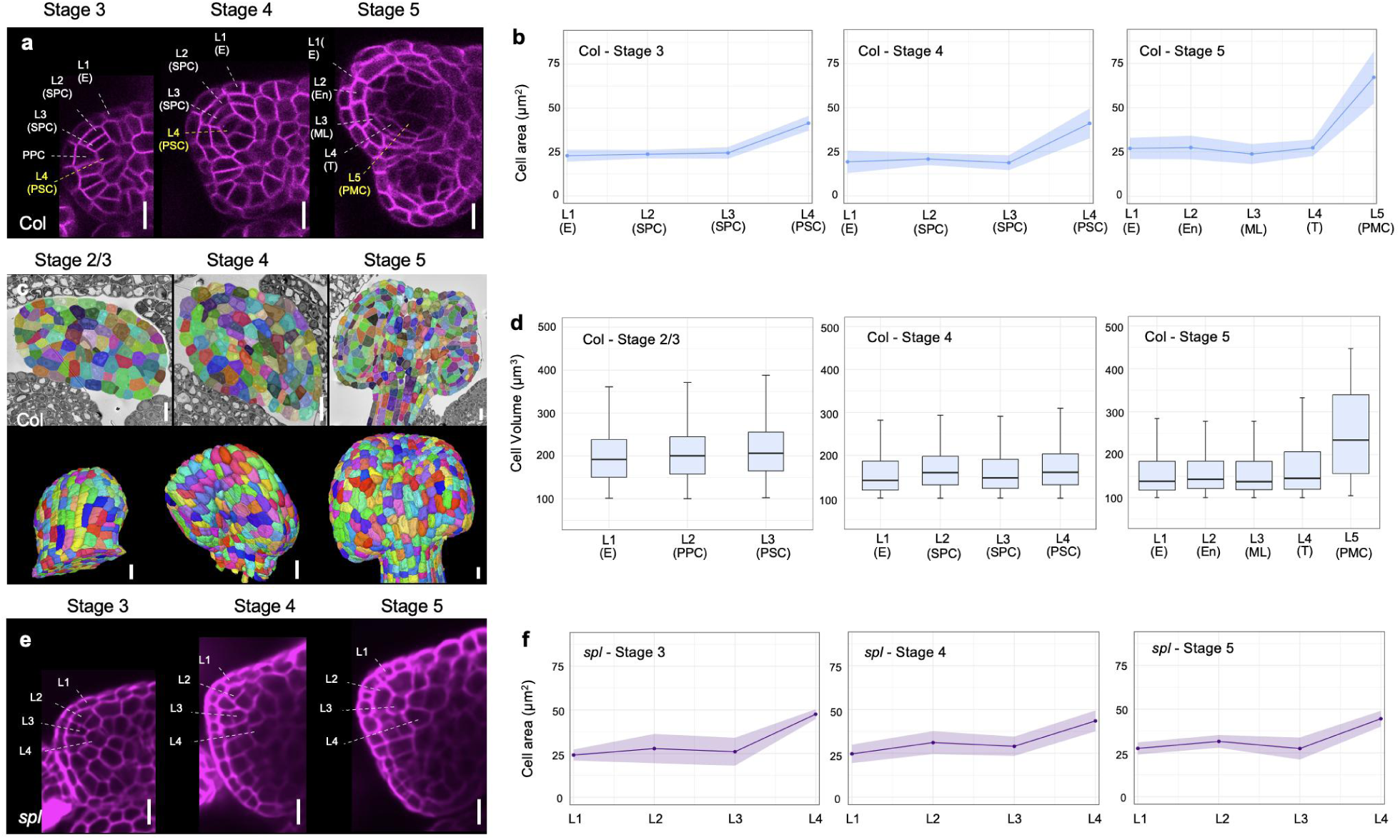
Cell size in WT and *spl* anthers. (a) Optical section of confocal images of WT anthers at stages 3-5, showing the enlargement of GC cells (marked with yellow letters). (b) Cell size in WT anther lobes quantified with cell area according to a (n = 4 anther lobes were analyzed for stage 3; n = 5 for stage 4; n = 7 for stage 5). (c) Display of the 3D reconstruction results for stage 2/3, stage 4 and stage 5 Arabidopsis anther serial electron microscope (EM) images. upper panel, showing the segmented cells in a single plane; lower panel, surface view of 3D reconstruction images. (d) Cell size in WT anther lobes quantified with cell volume according to c (n = 6 anther lobes were analyzed for stage 2/3; n = 6 for stage 4; n = 4 for stage 5). (e) Optical section of confocal images of *spl* anthers at stages 3-5. (f) Cell size in *spl* anther lobes quantified with cell area according to e (n = 4 for stage 3; n = 6 for stage 4; n = 6 for stage 5). Scale bars, 10 μm.

**Supplementary Fig. 3.**
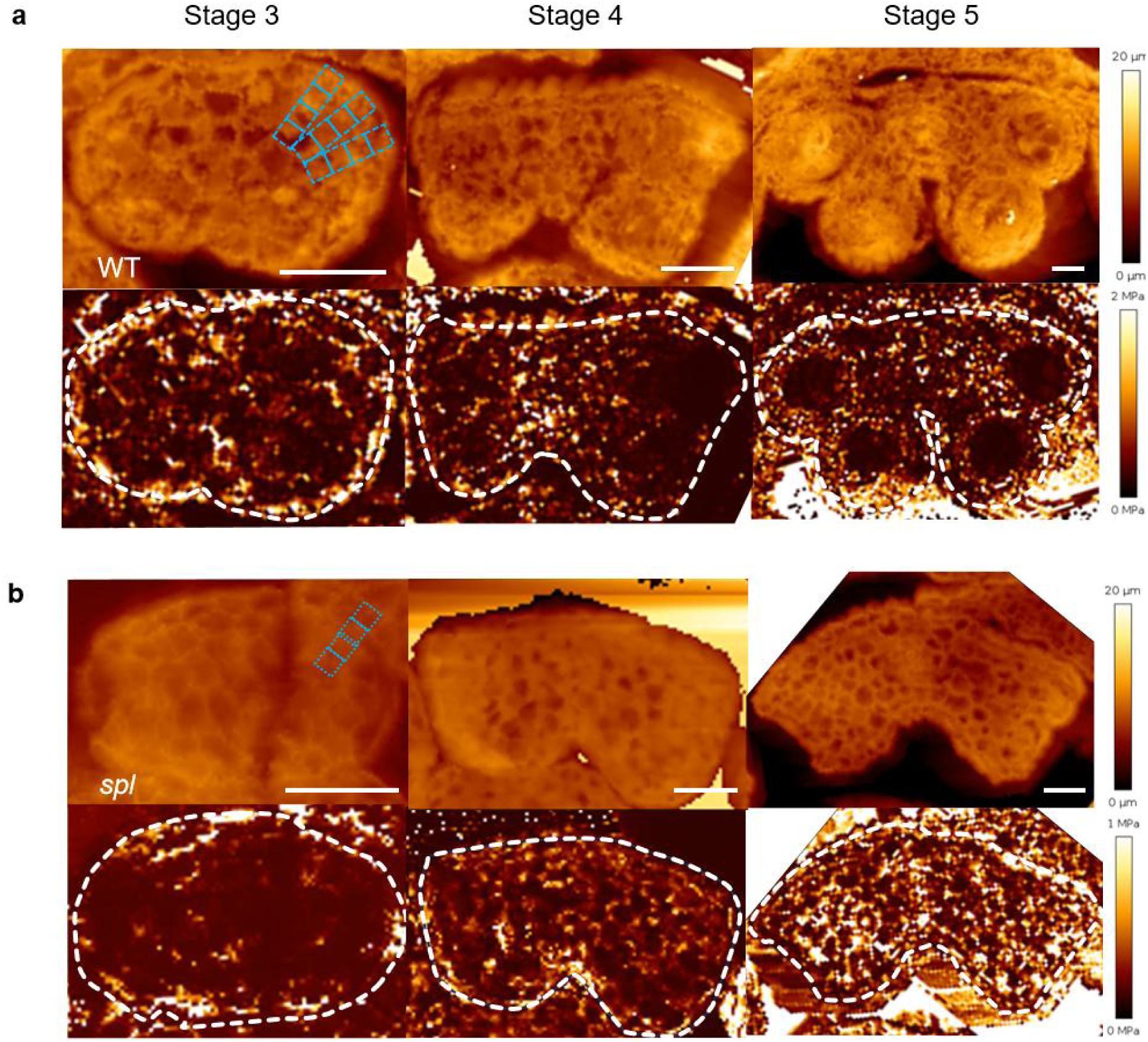
AFM measurement of anther stiffness during germ cell production. (a) Representative height (upper panel) and Young’s modulus maps (lower panel) on cross-sections of WT anthers at stages 3-5. Dashed blue boxes represent regions that were selected for quantification. The area for each blue box is about 5µm×5µm in size. Three radial-directed measurements along linear regions are collected for statistical analysis. Dashed white lines in the lower panel mark the boundaries of anther shape based on height map (upper panel). (b) Representative height (upper panel) and Young’s modulus maps (lower panel) of cross-sections of *spl* anther at stages 3-5, which are obtained by AFM. One of three radial-directed measurements along linear regions, taken into account for the quantifications, are shown. Scale bars, 20 μm.

**Supplementary Fig. 4.**
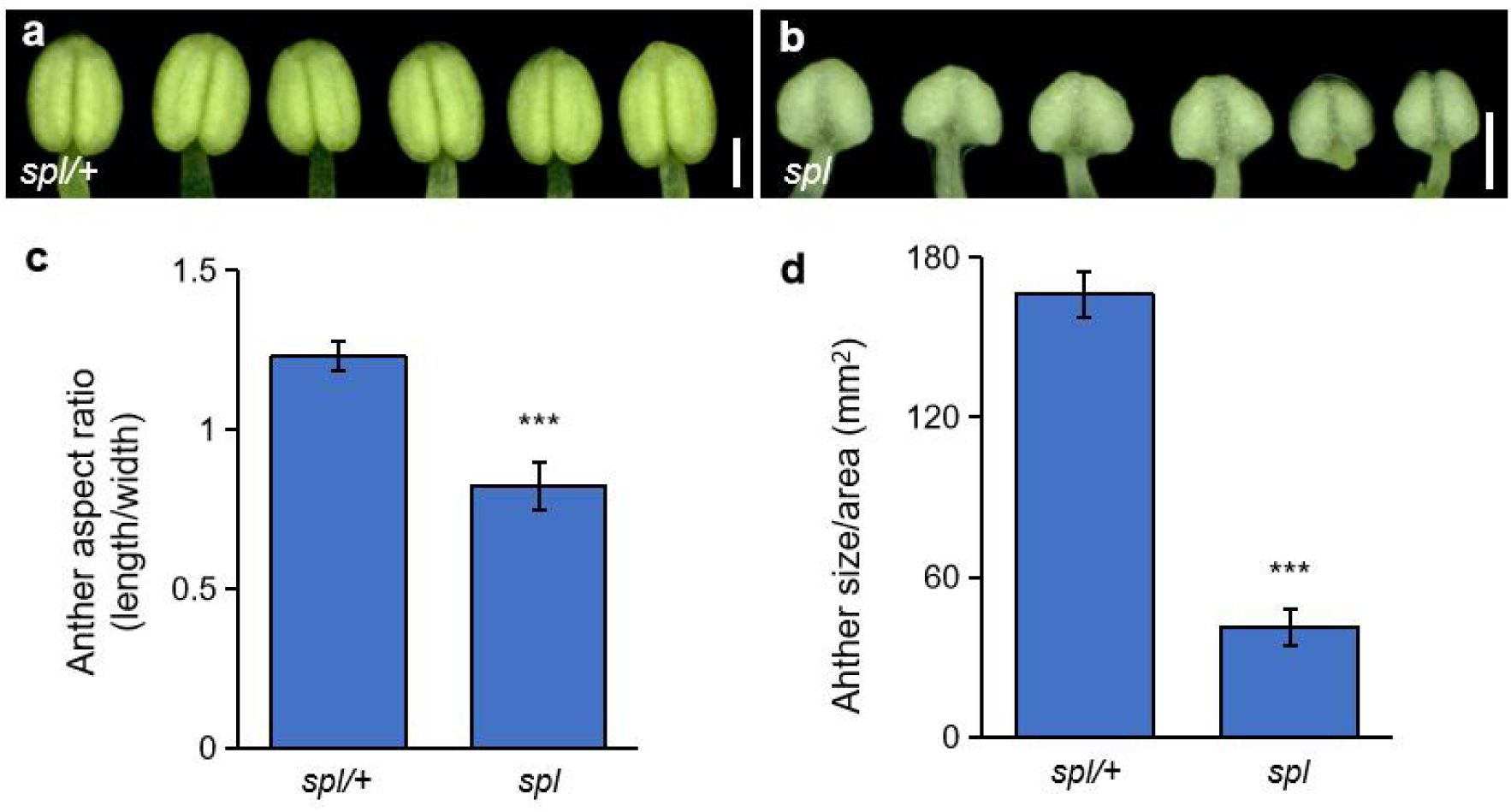
The anther shape and size of *spl* mutant. (a-b) Representative stamen (at stage 12) morphology of *spl/+* (a) and *spl* (b) collected from single flower bud respectively. (c-d) Quantification of anther aspect ratio (length/width) (c) and anther size/area (d) (n = 18 anthers were analyzed for each genotype). Values are mean ±SD, ***Student’s *t*-test *P* < 0.001. Scale bars, 250 µm.

**Supplementary Fig. 5.**
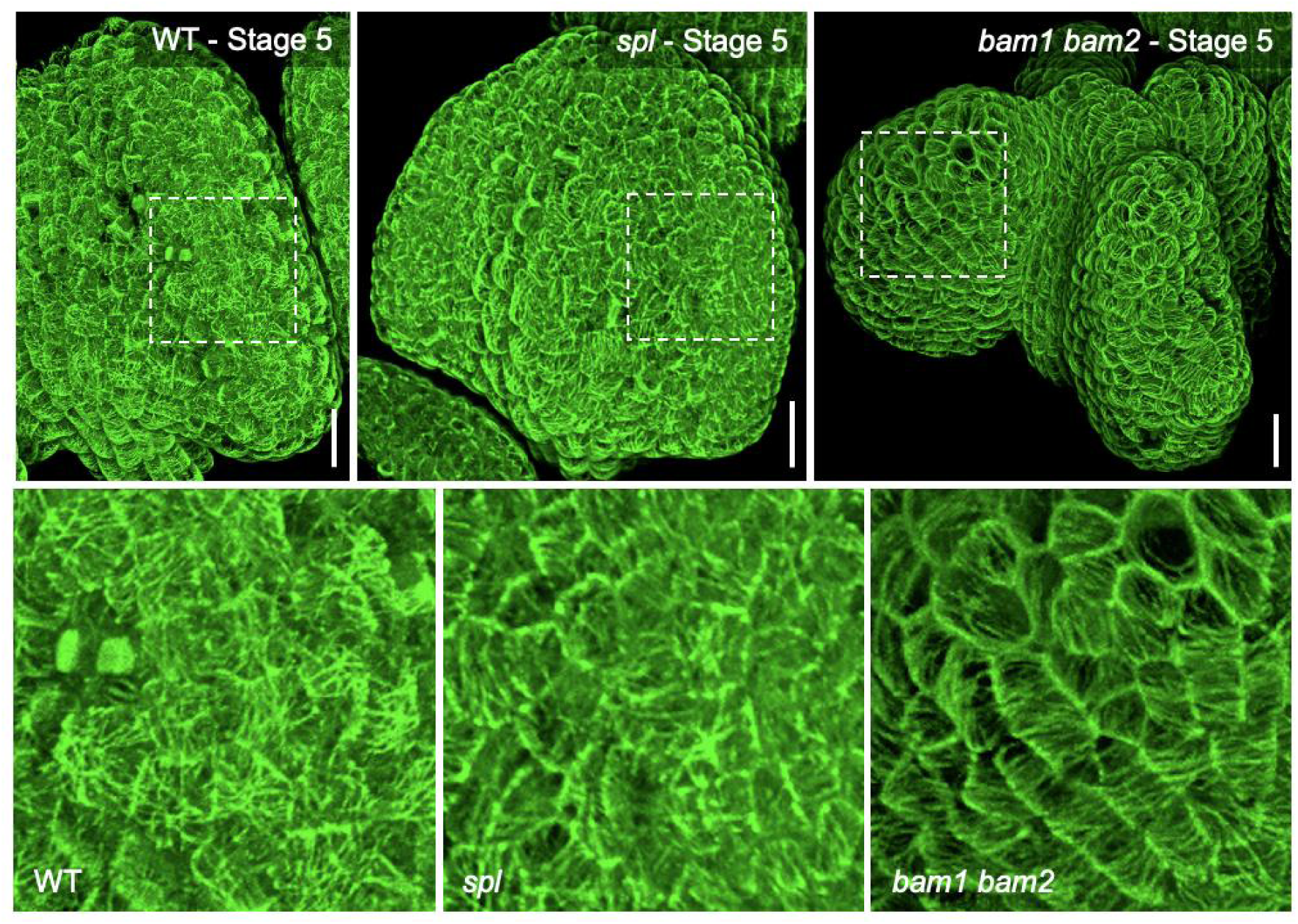
CMTs alignment on the surface of WT, *spl* and *bam1 bam2* anthers at stage 5. Upper panel, 3D reconstruction images showing the CMTs orientation on the anther surfaces. Bottom images are the magnified areas marked by a box in the images above. Scale bars, 10 µm.

**Supplementary Fig. 6.**
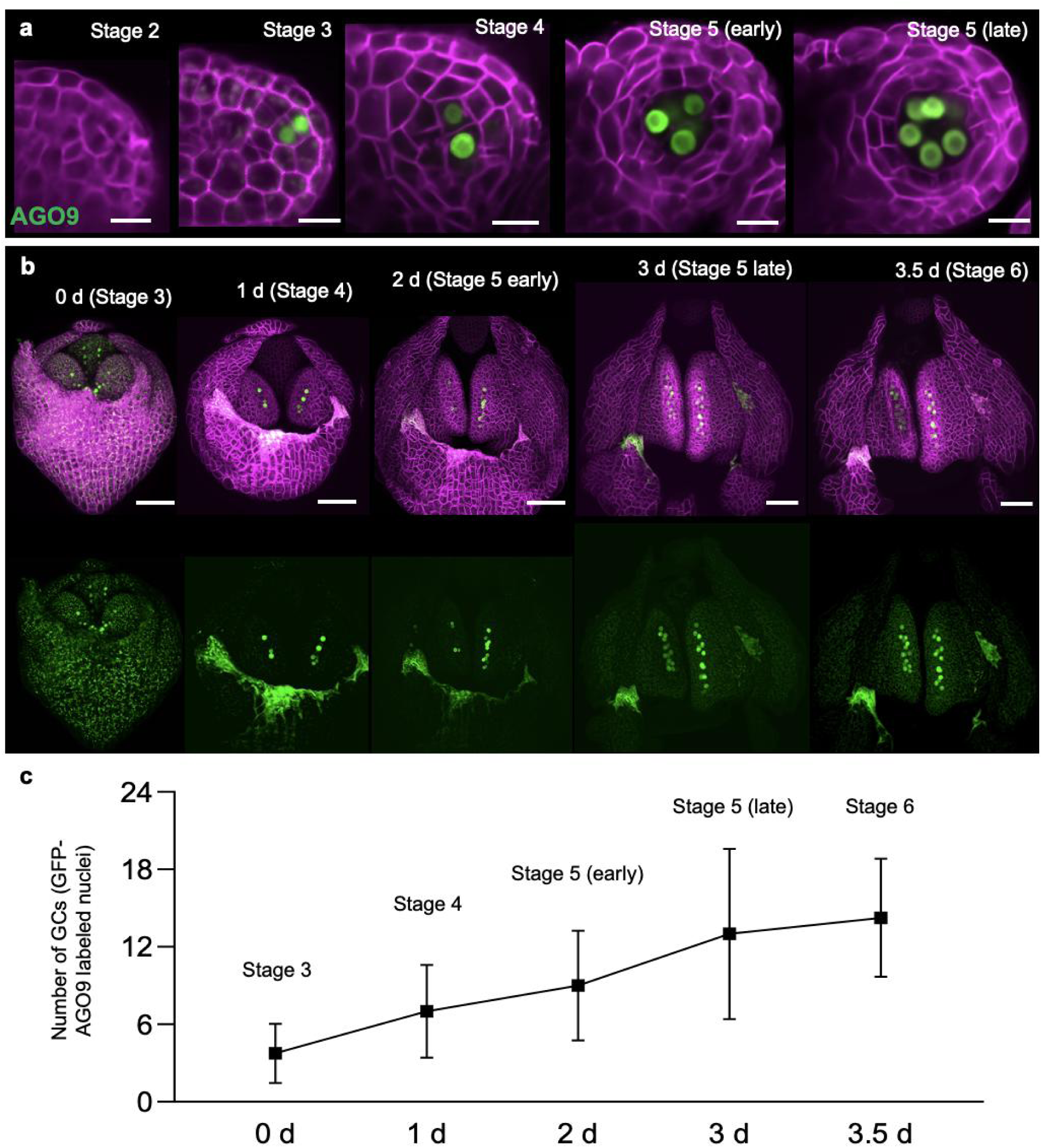
The expression pattern of GFP-AGO9 in developing anthers. (a) GFP-AGO9 expression pattern in different anther developmental stages. Stage 2-3, one theca; stage 4-5, one lobe. (b) 3D view of GFP-AGO9 distributions in anthers starting from stages 3 (0 d) to 6, which were grown on the ACM (apex culture medium) for 1 d, 2 d, 3 d and 3.5 d. Upper panel, merged images of plasma membrane marker LTI6b-tdTomato (magenta) and germ cell marker GFP-AGO9 (green). The lower panel, GFP channel alone. (c) Quantification of germ cell numbers, indicated by the number of GFP-AGO9 labeled nuclei, at different anther stages and growing days on the ACM. Scale bars, 10 µm (a), 50 µm (b).

**Supplementary Fig. 7.**
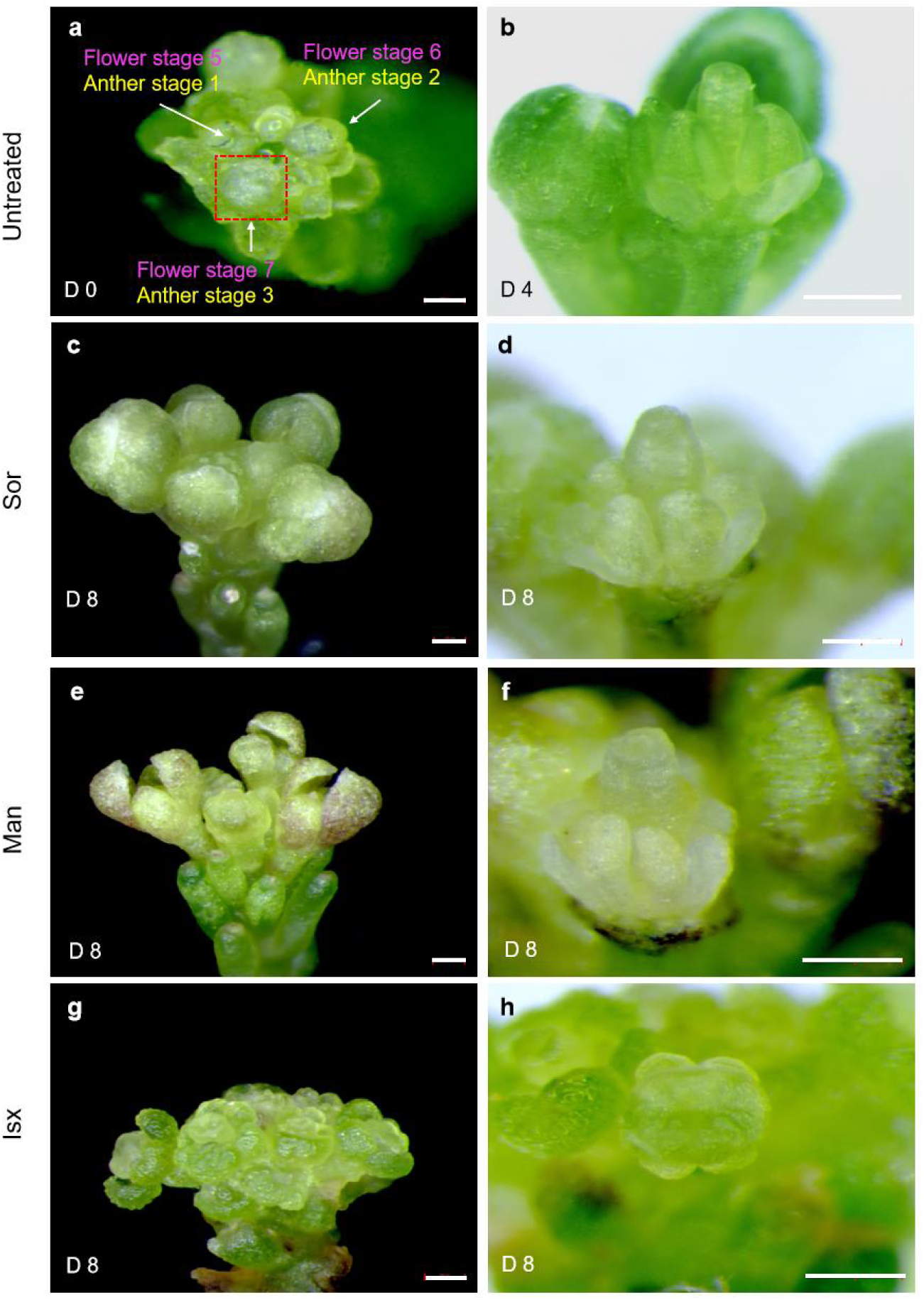
Phenotypic analysis of anthers under sorbitol, mannitol, and isoxaben treatments. (a-h) The inflorescence of LTI6b-tdTomato GFP-AGO9 plant w/o (a-b) or w/ treatment of sorbitol (c-d), mannitol (e-f) and isoxaben (g-h). D 0 (day 0) in (a) showing the appearance of the inflorescence before treatment. Three successive stages for the flowers and anthers are marked with magenta and yellow respectively. The dashed red frame marked out the oldest flower (at floral stage 7/anther stage 3) left on the inflorescence after the dissections. D 4 (day 4) in (b) represents untreated inflorescence grown on the ACM for 4 days. The sepals were dissected away to show the stamens. For the treatment groups, the right images (d, f, h) show the dissected flowers from the oldest ones on the inflorescence shown left (c, e, g) which are grown on the ACM for 8 days (D 8). Scale bars, 200 µm.

**Supplementary Fig. 8.**
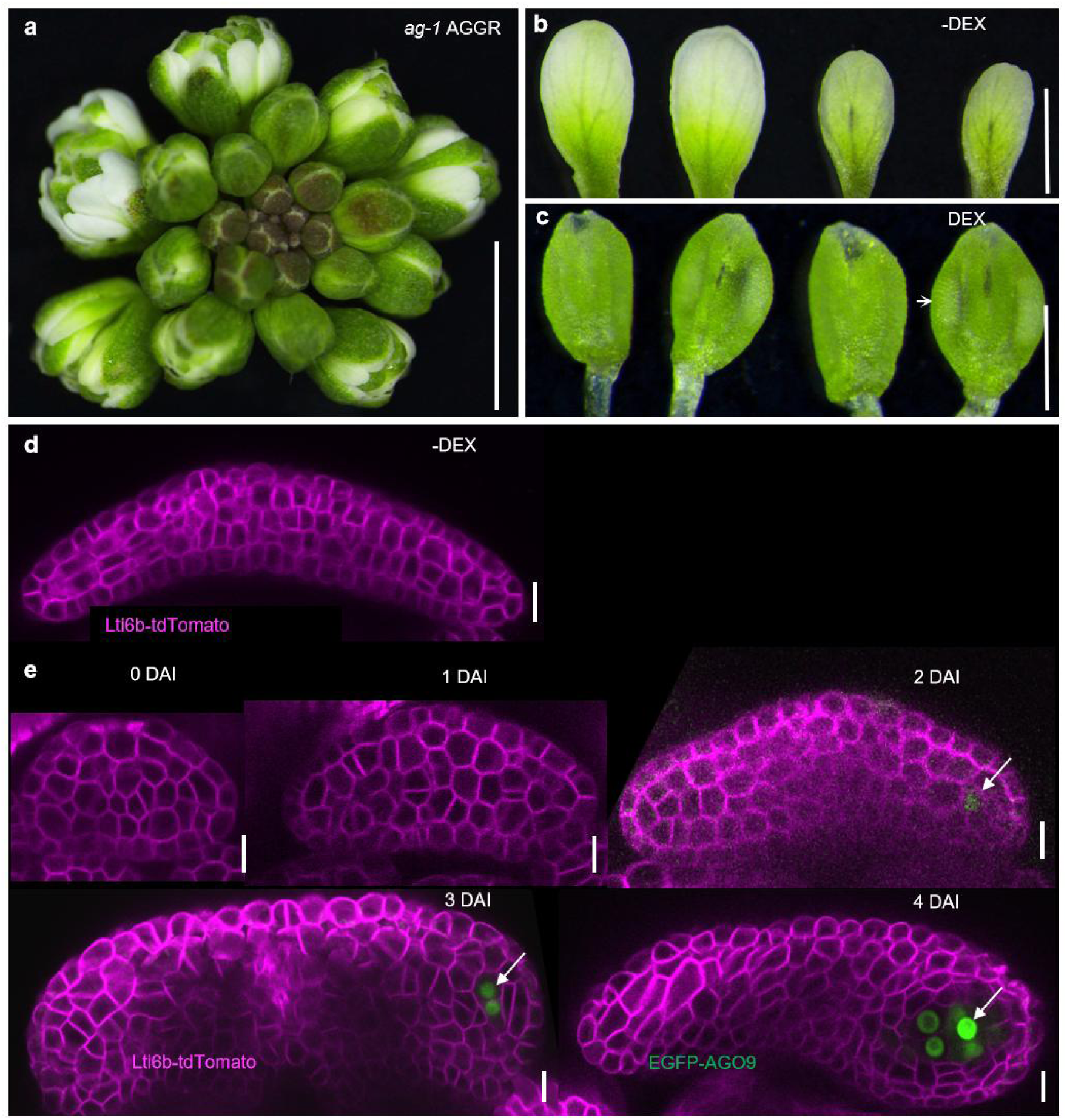
The GC induction in the petals of *ag-1* AGGR under DEX treatment. (a) The appearance of *ag-1* AGGR inflorescence. (b-c) Petals from *ag-1* AGGR inflorescence that growing on the ACM for 6 days without (b) or with (c) DEX treatment. White arrowhead indicates anther locule. (d) Optical cross-section view of one petal without DEX treatment. (e) The generation of GCs were tracked by GFP-AGO9 after single DEX treatment, arrows indicate the GC formation where GFP-AGO9 is activated. The plasma membrane is marked by LTI6b-tdTomato shown in magenta. Scale bars, 500 µm (a-c), 10 µm (d-e),

**Supplementary Fig. 9.**
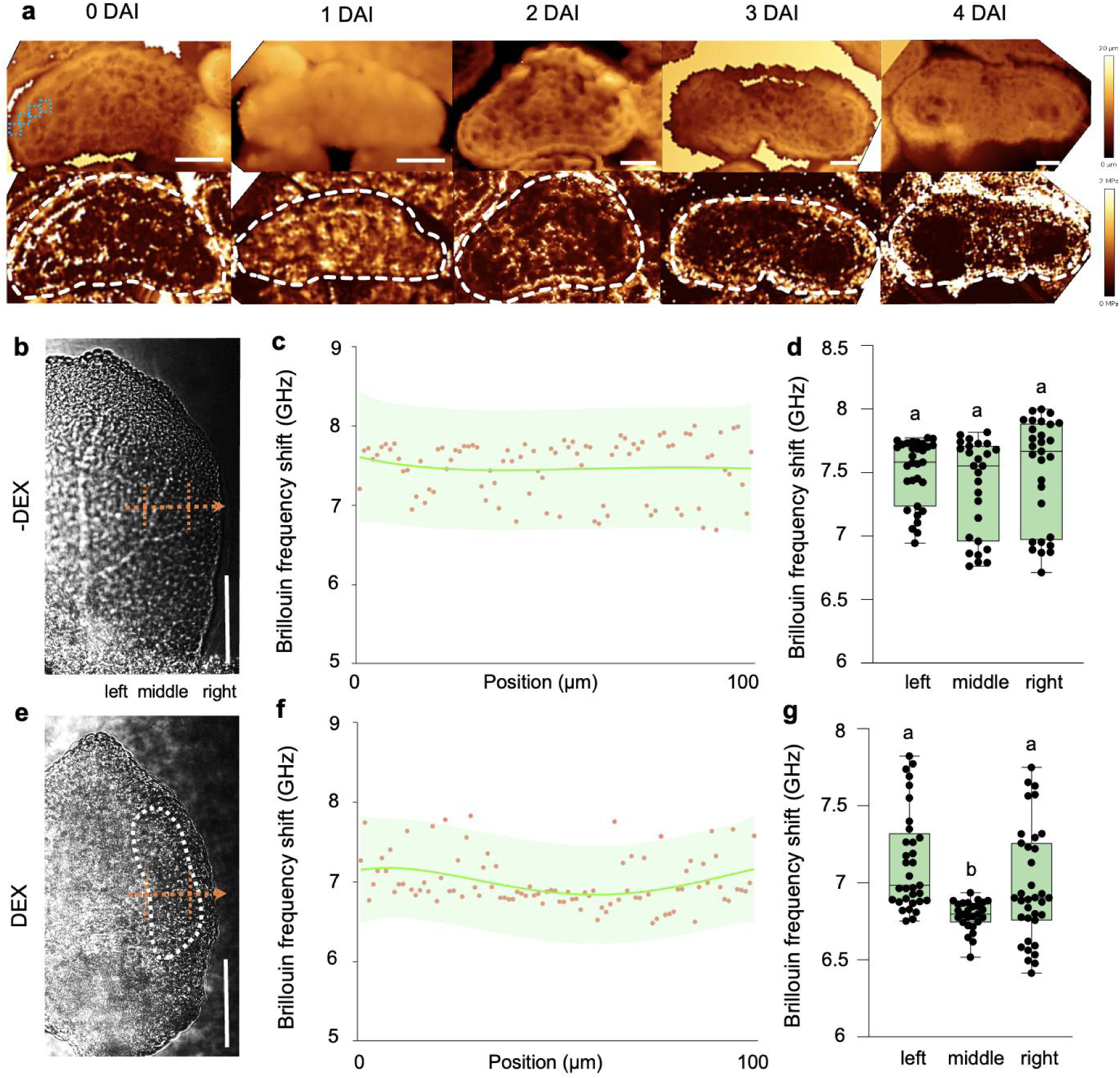
The CW stiffness patterning in *spl* anthers and *ag-1 AGGR* petals. Representative height (upper panel) and Young’s modulus maps (lower panel) of cross-sections of *ag-1 AGGR* petals at different growing days, that are obtained by AFM. Dashed blue boxes represent regions that were selected for quantifications. The area for each blue box is about 5µm×5µm in size. One of three radial-directed measurements along linear regions, taken into account for the quantifications, are shown. Dashed white lines mark the boundaries of petals based on the height map. (b-g) Tissue stiffness measured by Brillouin microscope. Brillouin frequency shift (GHz) was measured across the radial axis of half of petal that without (b-d) or with DEX (e-g) treatment by line scanning (about 100 µm, dashed orange arrows). Dashed white lines define the boundaries of Microsporocytes. Brillouin frequency shift measurements were performed after 6 days of treatment. Dots in (c and f) represent individual measurements, with lines representing corresponding regression curves with a 95% confidence interval (shadowed). The dashed vertical orange lines in (b and e) divide the 100 microns into three parts (left, middle, right), and dots fall in these areas are used for statistics. Values are mean ±SD. Different letters on graphs indicate significant difference (*P* < 0.05) calculated by one-way ANOVA followed by Tukey’s test. Scale bars, 20 µm (a). 100 µm (b, e).

**Supplementary Fig. 10.**
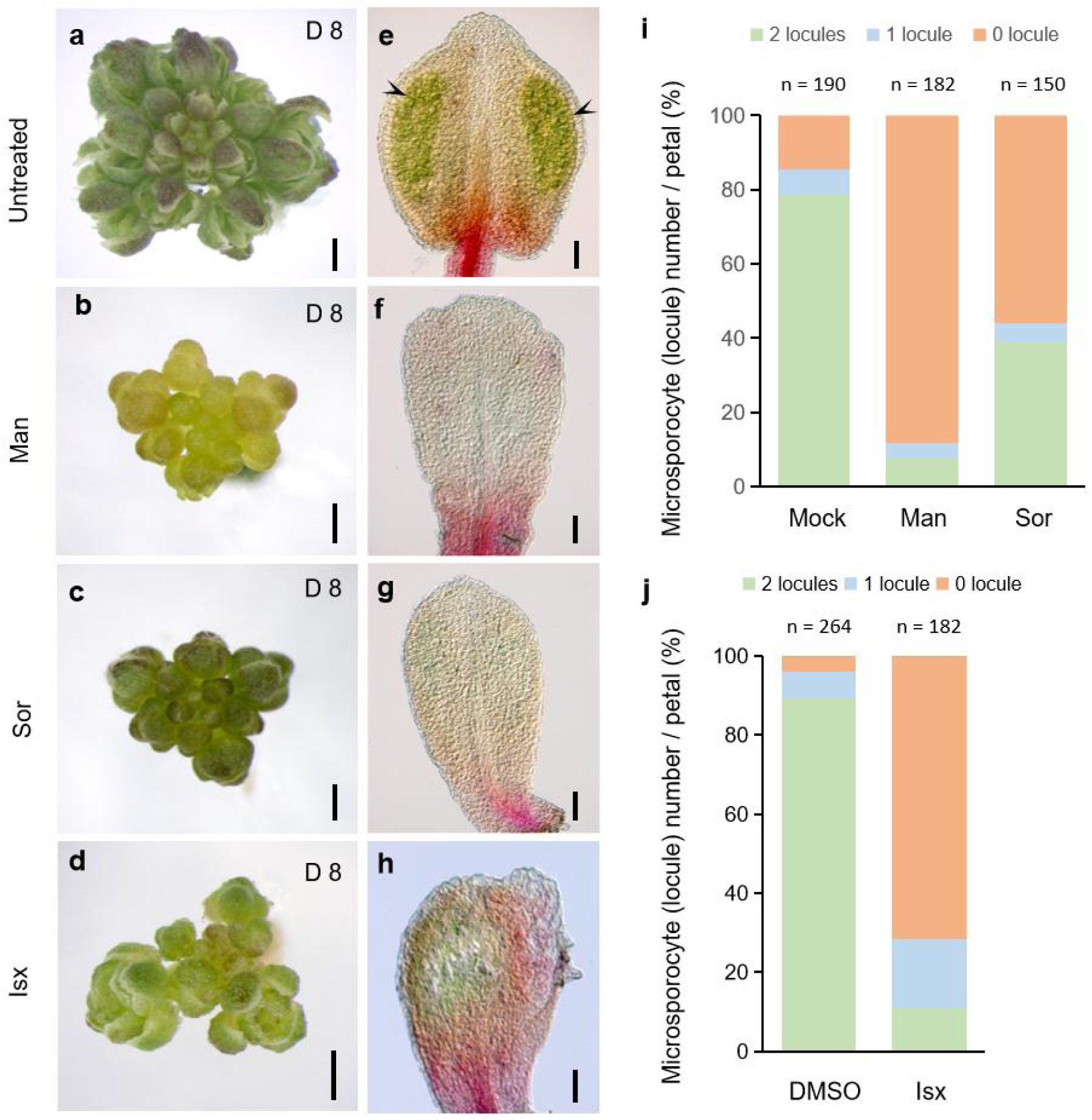
Germ cell production is affected by mechanical perturbations. (a-d) The phenotype of inflorescence of *ag-1* AGGR that untreated (a) or treated with mannitol (b), sorbitol (c) and isoxaben (d). (e-h) Alexander’s staining to indicate germ cell production in petals. Black arrowheads indicate the microspores. Note that in some extreme cases, after the mechanical perturbations, there is no GCs formation along the petal margins (f-h). (i-j) Quantitative analysis of the numbers of microsporocytes/locules (harboring GCs) formation at D 8 w/o treatment (Mock and DMSO) or w/ the treatments of mannitol, sorbitol (i) and isoxaben (j). Scale bars, 500 µm (a-d), 50 µm (e-h).

**Supplementary table 1.**
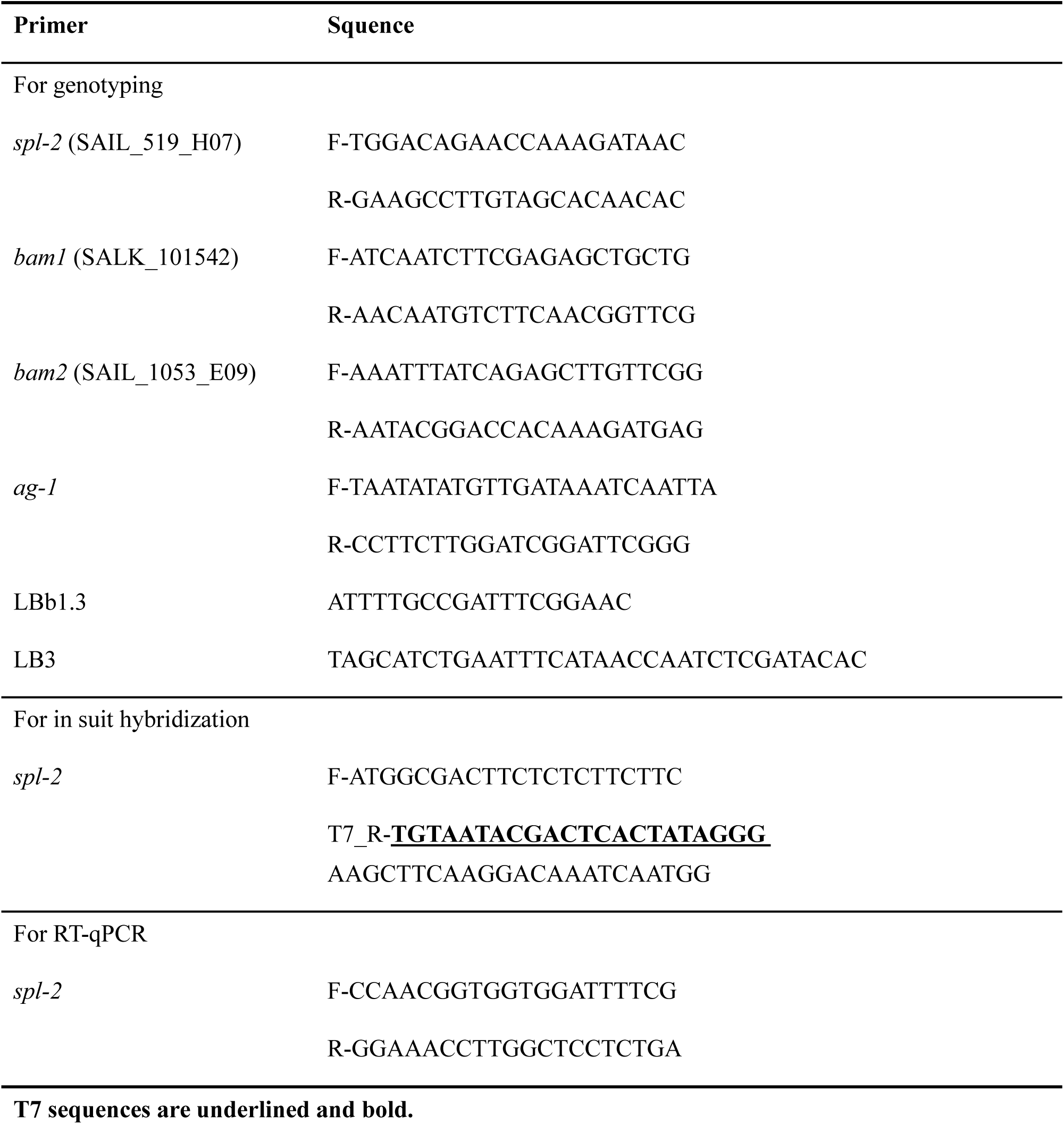
Primer sequences used in this study Primer Squence.

